# Label-free imaging of macrophage phenotypes and phagocytic activity in the human dermis *in vivo* using two-photon excited FLIM

**DOI:** 10.1101/2021.11.29.470361

**Authors:** M. Kröger, J. Scheffel, E. A. Shirshin, J. Schleusener, M. C. Meinke, J. Lademann, M. Maurer, M. E. Darvin

**Affiliations:** Charité – Universitätsmedizin Berlin, corporate member of Freie Universität Berlin, Humboldt-Universität zu Berlin, and Berlin Institute of Health, Department of Dermatology, Venerology and Allergology, Charitéplatz 1, 10117 Berlin, Germany; Lomonosov Moscow State University, Faculty of Physics, 119991, Leninskie gory 1/2, Moscow, Russia

**Author notes:** Corresponding author: Maxim E. Darvin.

## Abstract

Macrophages (MΦs) are important immune effector cells that promote (M1 MΦs) or inhibit (M2 MΦs) inflammation and are involved in numerous physiological and pathogenic immune responses. Their precise role and relevance, however, is not fully understood because of the lack of non-invasive quantification methods. Here, we show that two-photon excited fluorescence lifetime imaging (TPE-FLIM), a label-free non-invasive method, can visualize MΦs in human dermis *in vivo*. We demonstrate *in vitro* that human dermal MΦs exhibit specific TPE-FLIM properties that distinguish them from the main components of the extracellular matrix and other dermal cells. We visualized MΦs, their phenotypes and phagocytosis in the skin of healthy individuals *in vivo* using TPE-FLIM. Additionally, machine learning identified M1 and M2 MΦs with a sensitivity of 0.88±0.04 and 0.82±0.03 and a specificity of 0.89±0.03 and 0.90±0.03, respectively. In clinical research, TPE-FLIM can advance the understanding of the role of MΦs in health and disease.

## Introduction

Macrophages (MΦs) are important immune effector cells in organs and tissues that act as border junctions to environments such as the gut, the airways, and the skin (Elhelu, 1983). Skin MΦs (Dong et al., 2016; Estandarte et al., 2016; Ryter, 1985) originate from circulating monocytes (Geissmann et al., 2010; Gordon and Taylor, 2005) via the same infiltration route into the dermis as monocyte-derived dendritic cells (Schmid and Harris, 2014) (Fig. 1a) and are mainly located in the papillary and reticular dermis in close proximity to blood vessels (Weber-Matthiesen and Sterry, 1990) (Figs. 1b-d). It has been known for more than 30 years that skin MΦs are abundant and heterogeneous, based on their morphology, localization and staining properties (Weber-Matthiesen and Sterry, 1990). More recently, skin MΦs have been classified based on their function, and they fall into two phenotypes referred to as inflammation-promoting M1 MΦs (classically activated) and anti-inflammatory M2 MΦs (alternatively activated) (Duque and Descoteaux, 2014) (Fig. 1a). M1 MΦs are activated by viral and bacterial infection (Benoit et al., 2008; Ferrer et al., 2019; Malmgaard et al., 2004), interferon-γ, lipopolysaccharide (LPS), and tumor necrosis factor (TNF), which is known as the classical activation pathway (Li and Liu, 2018). M2 MΦs are alternatively activated in response to IL-4, IL-13 and IL-33 (Furukawa et al., 2017; Sica and Mantovani, 2012).

**Figure 1.**
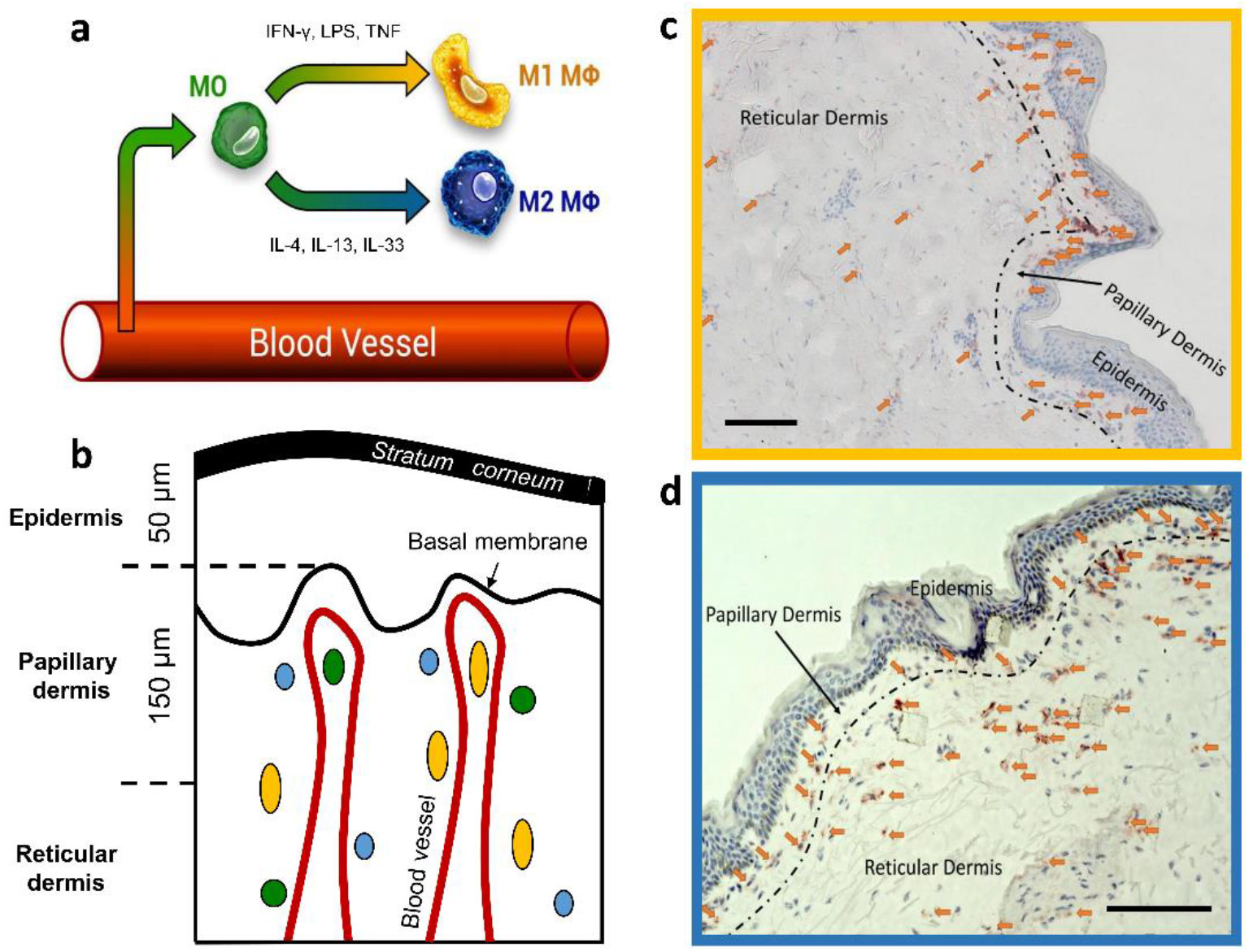
Dermal monocyte skin infiltration and CD68 stained M1 and CD163 stained M2 MΦs distribution in excised human skin. Schematic illustration of monocyte (MO) (green) infiltration into tissues and macrophage (MΦ)-polarization into M1 MΦs (yellow) via IFN-γ, LPS and TNF and M2 MΦs (blue) via IL-4, IL-13 and IL-33 (a). Schematic of skin with exemplary locations of monocytes (green), M1 MΦs (yellow) and M2 MΦs (blue) (b). Density of M1 MΦs (marked with arrows) stained with CD68 (c) and M2 MΦs (marked with arrows) stained with CD163 (d) in 10 μm thick cryo-section. Scale bar 100 μm. Recently, this paradigm has been challenged, as many signals modulate MΦ functions, resulting in a continuum of states between the M1 and M2 phenotypes(Mendoza-Coronel and Ortega, 2017; Murray et al., 2014). However, for simplicity, the terms M1 and M2 MΦs are used here.

Skin M1 MΦs are held to contribute to dermal innate immunity and homeostasis. This is supported by reports that M1 MΦs can phagocyte objects up to 20 μm in size (Morhenn et al., 2002), promote skin inflammatory and immune responses (Remmerie and Scott, 2018; Theret et al., 2019; Yanez et al., 2017), and produce nitric oxide and other reactive oxygen species (ROS) (Forman and Torres, 2001; Rendra et al., 2019). Skin M2 MΦs, on the other hand, are thought to promote dermal repair, healing and regeneration, for example by contributing to the formation of the extracellular matrix (Ploeger et al., 2013).

The precise role of skin MΦs and their M1 and M2 phenotypes in health and disease remains to be elucidated. Efforts to do so include their quantification in healthy human skin and in lesional and non-lesional skin of patients with skin diseases. Currently, the most common approach is to obtain skin biopsies and to visualise MΦs by immunohistochemistry. Skin biopsies, however, come with several important limitations, which include scarring, the risk of infection and bleeding, and artificial findings caused by the use of local anesthesia. Also, histopathological analyses of skin biopsies are not well suited for characterizing MΦ functions such as phagocytosis and for long term monitoring of MΦ distribution in the skin. Fluorescence lifetime imaging (FLIM) employs NAD(P)H and fluorescence decay parameters of cellular compartments as specific indicators of cell types and phenotypes (Alfonso-García et al., 2016; Heaster et al., 2021). Combined with two-photon tomography, two-photon excited fluorescence lifetime imaging (TPE-FLIM) allows for label-free and non-invasive imaging of dermal cells. For instance, TPE-FLIM allows for *in vitro* imaging of mast cells, fibroblasts, neutrophils, and dendritic cells and *in vivo* imaging of mast cells in human skin (Kröger et al., 2020). Whether or not TPE-FLIM can be used to visualise human skin MΦs, their M1 and M2 phenotypes, and their functions, is currently unknown. There are, however, several independent lines of evidence that support this approach: First, previous studies have shown that TPE-FLIM can distinguish MΦs from other dermal cells and extracellular matrix (ECM), without prior labelling (Kröger et al., 2020). Second, the capillaries of the papillary dermis, which often are in a close proximity with MΦs, show distinct TPE-FLIM signatures and are readily visualised (Shirshin et al., 2017). Third, M1 and M2 MΦs come with unique cytokine patterns, and the TPE-FLIM signatures of these cytokines and patterns could help to tell the two phenotypes apart. Finally, TPE-FLIM can distinguish between functional states of dermal cells, e.g. resting and activated mast cells *in vivo*, MΦs *ex vivo* (Kröger et al., 2020) and T-cell activation *in vitro* (Walsh et al., 2021) may, therefore, potentially allow for monitoring MΦ functions *in vivo* (Szulczewski et al., 2016). Taken together, the morphological features of skin MΦs, their localization in the skin, and the expected differences in fluorescence decay parameters between MΦ phenotypes as well as between other dermal cells, make TPE-FLIM a promising strategy for their detection (Yakimov et al., 2019).

Here, we first investigated human skin MΦs, *in vitro*, for their TPE-FLIM properties and how these differ from those of the main components of the ECM and other dermal cells such as fibroblasts, mast cells, and dendritic cells. We then applied the identified MΦ TPE-FLIM signatures to investigate MΦs and their phenotypes in human skin biopsies, combined with traditional immunohistochemistry-based visualization. Finally, we used TPE-FLIM *in vivo* in humans to study skin MΦs, their phenotypes and functions, and we developed, tested, and characterised TPE-FLIM signature-based machine learning algorithms for the detection of skin MΦs.

## Results

### In vitro monocyte-derived M1 and M2 MΦs show distinct TPE-FLIM parameters

The TPE-FLIM images of monocytes isolated from human peripheral blood mononuclear cells (PBMC) showed a round morphology (diameter of up to 10 μm) with a barely visible nucleus, homogeneously distributed cell content, and regular borders with no membrane extensions (Fig. S1). MΦs differentiated from PBMC and polarised towards M1 MΦs with interferon-γ (IFN-γ; *n*=21) and towards M2 MΦs with interleukin-4 (IL-4; *n*=27) were similar in size, ranging from 10–12 μm (Fig. 2a, 2b and 2c). M1 and M2 MΦs showed comparable overall TPE-AF intensities, but they differed significantly in several other features. M1 MΦs also showed numerous bright spots (typical size is 2–3 μm), likely vacuoles and mitochondria, had less visible borders, and exhibited higher TPE-AF intensity than M2 MΦs. In contrast, M2 MΦs were characterised by distinct borders with filopodia (Fig. 2b; Fig. S2), which were rarely seen in M1 MΦs (Fig. 2a).

**Figure 2.**
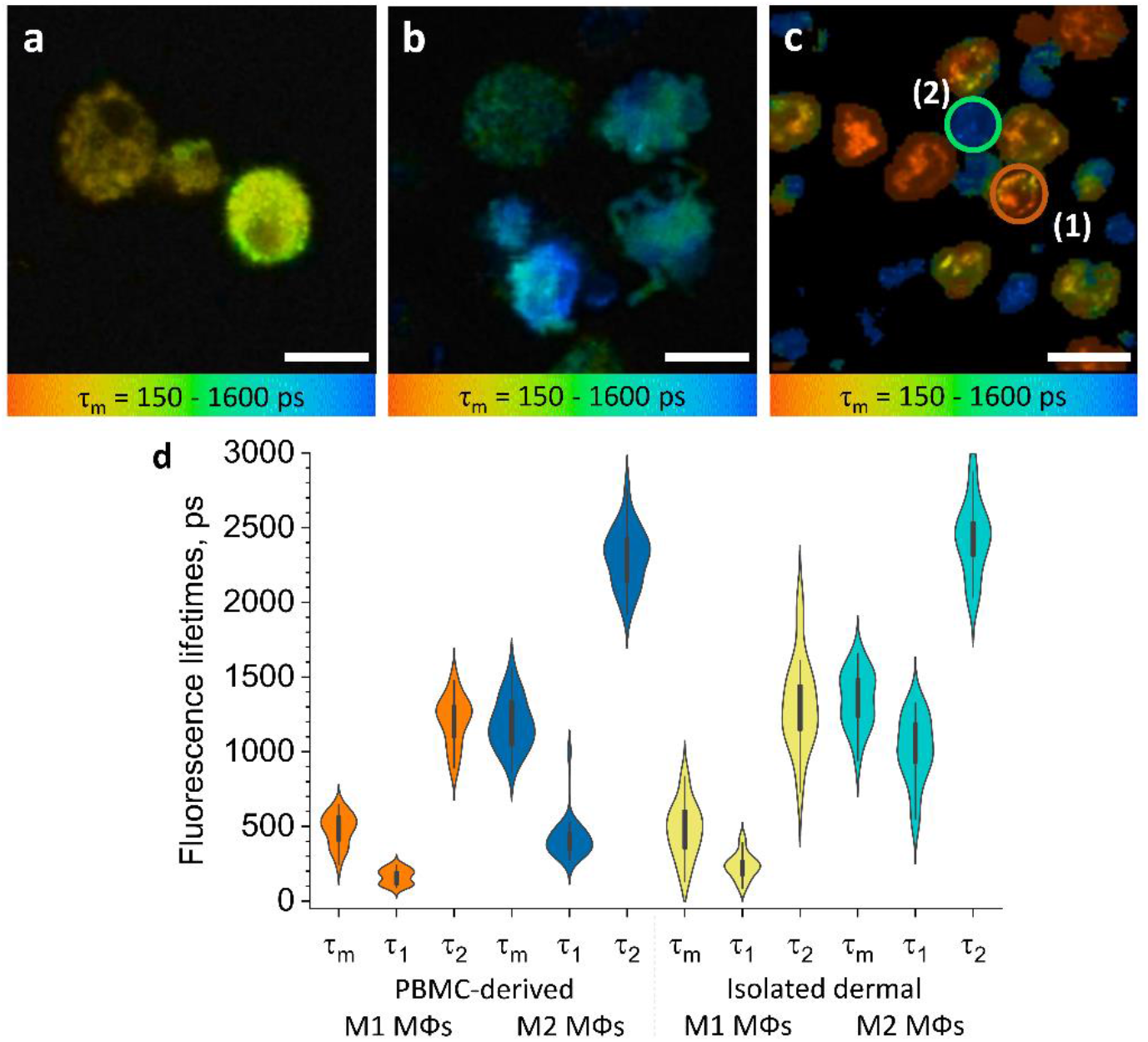
MΦs polarised from PBMC and isolated dermal MΦs show distinct TPE-FLIM signatures. TPE-FLIM *τ_m_* images (mean fluorescence lifetime *τ_m_* in the 150–1600 ps range) of monocyte-derived M1-polarised (IFN-γ) MΦs (a), monocyte-derived M2-polarised (IL-4) MΦs (b), and isolated human dermal M1 MΦs (1) and M2 MΦs (2) (c). Scale bar: 10 μm. The distribution of TPE-FLIM parameters *τ_1_*, *τ_2_* and *τ_m_* for monocyte-derived M1-polarised MΦs (*n*=21, orange), M2-polarised MΦs (*n*=27, dark blue), and isolated dermal M1 MΦs (*n*=34, yellow), M2 MΦs (*n*=28, light blue) (d). The boxplot represents 25–75% of the values.

M1 and M2 MΦs also differed in their TPE-FLIM parameters *τ_1_*, *τ_2_*, and *τ_m_* (Fig. 2d). TPE-AF decay times were significantly shorter in M1 MΦs (*n*=21) than in M2 MΦs (*n*=27; *p*<0.05), and both MΦs differed significantly, in their TPE-FLIM parameters, from monocytes (*n*=15; *p*<0.05; Table 1).

**Table 1.** TPE-FLIM parameters for for investigated dermal and epidermal cells. TPE-FLIM parameters *τ*_1_, *τ*_2_, *τ*_m_, *a_1_/a_2_* and TPE-AF intensity of monocyte-derived M1 and M2 MΦs; dermal M1 and M2 MΦs isolated from the skin measured *in vitro*; M1 (CD68) and M2 (CD163) MΦs measured *ex vivo* in human skin cryo-sections; M1 and M2 MΦs observed on the forearm of healthy volunteers *in vivo*; monocytes; resting and activated human skin mast cells; dendritic cells; fibroblasts and neutrophils *in vitro*.

| | | Number<br>of cells | $\tau_m$<br>in ps | $\tau_1$<br>in ps | $\tau_2$<br>in ps | $a_1/a_2$ | TPE-AF<br>intensity,<br>photons /mW |
| --- | --- | --- | --- | --- | --- | --- | --- |
| *in vitro* | Monocyte-derived M1-polarised MΦs | 21 | 479±106 | 163±50 | 1,209±161 | 2.4±0.6 | 600±100 |
| *in vitro* | Monocyte-derived M2-polarised MΦs | 27 | 1,185±170 | 417±134 | 2,305±194 | 2.3±0.5 | 500±100 |
| *in vitro* | M1 isolated dermal MΦs | 34 | 461±175 | 225±84 | 1,289±278 | 4.8±3.4 | 3,000±500 |
| *in vitro* | M2 isolated dermal MΦs | 28 | 1,281±155 | 807±250 | 2,352±229 | 2.2±1.1 | 800±200 |
| *ex vivo* | **M1 MΦs (CD68)** | 8 | 458±50 | 190±38 | 1,504±133 | 4.1±0.7 | 3,000±500 |
| *ex vivo* | **M2 MΦs (CD163)** | 12 | 1,369±201 | 498±129 | 2,267±155 | 1.1±0.4 | 700±300 |
| *in vivo* | **M1 MΦs** | 35 | 477±105 | 196±40 | 1,698±172 | 5.0±2.8 | 686±165 |
| *in vivo* | **Phagocytosing M1 MΦs** | 2 | 195±44 | 105±10 | 1,272±89 | 14.7±4.5 | 1,100±150 |
| *in vivo* | **M2 MΦs** | 25 | 1,407±60 | 442±54 | 2,458±90 | 1.2±0.2 | 360±155 |
| *in vitro* | **PBMC-derived monocytes** | 15 | 989±111 | 491±130 | 2,025±301 | 1.8±0.5 | 700±130 |
| *in vitro* | **Resting mast cells** | 43 | 1,248±287 | 533±266 | 2,289±317 | 1.5±0.5 | 1,300±400 |
| *in vitro* | **Activated mast cells** | 13 | 862±268 | 288±130 | 1,920±287 | 2.5±2.0 | 900±200 |
| *in vitro* | **Dendritic cells** | 14 | 1,265±180 | 434±188 | 2,578±328 | 1.6±0.2 | 538±258 |
| *in vitro* | **Fibroblasts** | 6 | 921±81 | 429±51 | 1,983±137 | 0.5±0.1 | 469±137 |
| *in vitro* | **Neutrophils** | 21 | 1,074±109 | 714±250 | 1,795±600 | 1.5±0.5 | 500±115 |

IgG stimulation of IgG immune complex-sensitised M1 and M2 MΦs resulted in the release of inflammatory mediators, but did not lead to significant changes or reveal additional differences in TPE-FLIM parameters 2 and 5 days after differentiation of PBMC into MΦs (data not shown). Taken together, these findings indicate that monocyte-derived M1 MΦs and M2 MΦs can be identified and distinguished *in vitro* by their distinct TPE-FLIM signatures.

### MΦs isolated from periocular skin show TPE-FLIM parameters that are similar to those of in vitro monocyte-derived M1 MΦs or M2 MΦs

Human MΦs isolated from periocular dermal tissue and analysed by immunohistochemistry were irregularly shaped, with poorly defined borders, 8–10 μm in size, pericentral nuclei of 5–6 μm diameter with low fluorescence intensity, heterogeneously and irregularly distributed cellular content, and they exhibited a bright fluorescence multi-vacuolated cytoplasm with ≈1 μm diameter small bright spots, presumably related to mitochondria and/or vacuoles (Fig. 2c). Based on their TPE-FLIM parameters, dermal MΦs fell into two significantly different groups (Fig. 2d): group 1 (*n*=34), with stronger TPE-AF intensity (≈3,000±500 photons/mW) and shorter lifetimes, and group 2 (*n*=28), with a weaker TPE-AF intensity (≈800±200 photons/mW) with longer lifetimes (Fig. 2c; Fig. S3). The profiles of dermal MΦs in group 1 and 2 were similar to those of monocyte-derived M1 MΦs and M2 MΦs, respectively (Fig. 2d; Table 1). The biggest differences between group 1 / M1 MΦs and group 2 / M2 MΦs were shorter *τ_1_* and *τ_m_* as well as larger size (10.9±0.6 μm) in the former as compared to the latter (9.8±1.2 μm; *p*<0.05; Fig. 2c and 2d; Table 1). This suggests that human skin MΦs, based on their *in vitro* TPE-FLIM signatures, can be assigned to one of two phenotypes, where the first is similar to that of monocyte-derived M1 MΦs and the second is similar to that of monocyte-derived M2 MΦs.

### TPE-FLIM can distinguish between MΦs and other cells

To prove that the recorded TPE-FLIM signatures are unique for M1- and M2-polarized MΦs (Fig. 2), we performed TPE-FLIM measurements of other dermal cells *in vitro*, such as mast cells, dendritic cells, fibroblasts, monocytes and neutrophils. Their TPE-FLIM parameters, summarised in Table 1, are markedly different from those of the established signatures of M1- and M2-polarized MΦs. Thus, in addition to size, morphology, and internal vacuole structure, M1- and M2-polarized MΦs can be distinguished, from each other and other cells, by distinct TPE-FLIM parameters, a prerequisite for the visualization of skin MΦs *ex vivo* and *in vivo*. Table 1 is an extension of the results shown in Kröger *et al*.(Kröger et al., 2020)

### Immunohistochemistry confirms TPE-FLIM detection of M1 and M2 MΦs in human skin ex vivo

To test if the TPE-FLIM signatures established *in vitro* identify M1- and M2-polarized MΦs in human skin, we sequentially analyzed dermal biopsies by TPE-FLIM and conventional immunohistochemistry. The application of *in vitro* MΦ signatures to TPE-FLIM analyses of 13 human skin biopsy cryo-sections identified two distinct cell populations: The first showed a short mean fluorescence lifetime *τ*_m_ and high TPE-AF intensity, a feature of M1 MΦs (Table 1, Fig. 2d, Fig. 3a); the second population showed longer *τ*_m_ and significantly lower TPE-AF intensity, typical for M2 MΦs (Table 1, Fig. 2d, Fig. 3c). CD68-staining for M1 MΦs and CD163-staining for M2 MΦs confirmed that short *τ*_m_ cells with high TPE-AF intensity were, indeed, M1 MΦs (Fig. 3b) and that cells with longer *τ*_1_, *τ*_2_, *τ*_m_ with low TPE-AF intensity were, indeed, M2 MΦs (Fig. 3d, Fig. S4, Table 1).

**Figure 3.**
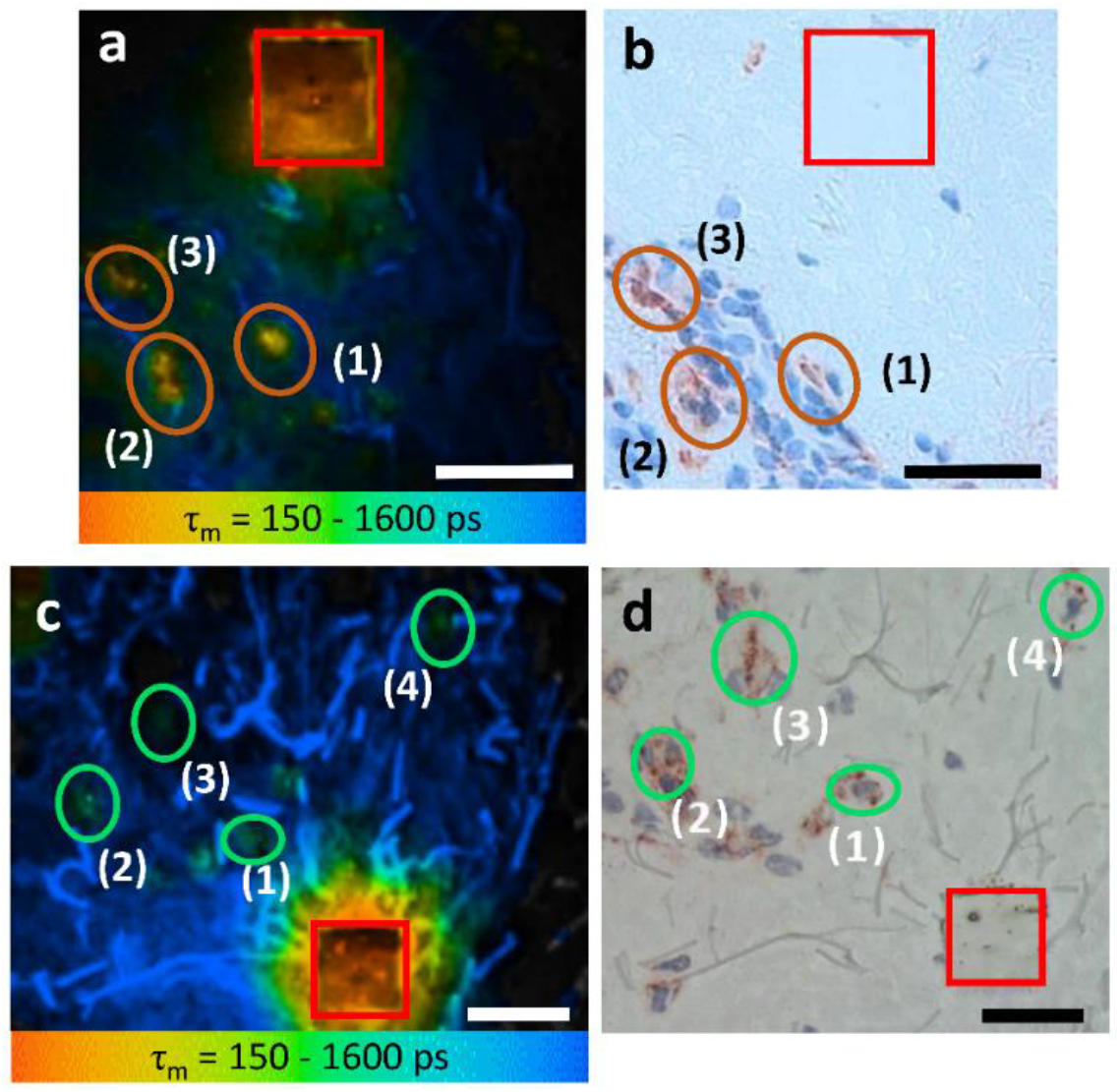
M1 and M2 MΦs *ex vivo* verified using TPE-FLIM parameters and immunohistochemistry-based bright field microscopy. Side by side comparison of TPE-FLIM τ_m_ image (mean fluorescence lifetime τ_m_ in the 150–1,600 ps range), which were measured label-free and then stained with CD68-antibody for M1 MΦs (a), and CDl63 - antibody for M2 MΦs (c) and corresponding bright field microscopic images (b) and (d). The excitation wavelength is 760 nm and laser power is 4 mW (a) and 2 mW (c). The M1 and M2 MΦs are marked with ellipses in (a,b) and in (c,d), respectively. The laser-burned labels (28 μm×28 μm) are marked in red. The suspected (a, c) and staining-proved (b, d) MΦs are marked with number (1, 2, 3 and 4). More M2 MΦs are observed in (d) compared to (c) due to the staining and visualization of the entire biopsy volume in (d) and limited imaging plane of the two-photon tomograph (1.2–2.0 μm) in (c). Images have been rotated and zoomed to match their orientation and size. Scale bar: 30 μm.

CD68-positive dermal M1 MΦs showed a heterogeneous appearance, ranging from flat and spindle shaped vessel lining to big intravascular with irregular borders and an irregular nucleus (Fig. 3b). The TPE-FLIM image of CD163-positive M2 MΦs show round to elliptical shaped cells with a significantly lower TPE-AF intensity (Fig. 3d). Of nine cells with a TPE-FLIM M1 MΦ signature, eight cells stained positive for CD68, and all CD68-positive cells had a TPE-FLIM M1 M12 of 14) were CD163-positive, and all CD163-positive cells had a TPE-FLIM M2 MΦ signature.

### TPE-FLIM visualises human skin M1 and M2 MΦs in vivo

Next, we used TPE-FLIM to assess the skin of 25 healthy individuals *in vivo*, and we identified and further characterised 35 and 25 MΦs with a M1 and M2 TPE-FLIM signature, respectively. *In vivo*, similar to biopsy sections, M1 and M2 MΦs were located in the papillary and reticular dermis at >80 μm depth (Fig. 4) and showed a density of >100 MΦs/mm^2^ (Figs. 1c, 1d). M1 MΦs fell into three distinct groups and were either flat and spindle shaped (Fig. 4a), slightly dendritic (Fig. 4b) or large and intervascular (Fig. 4c). M2 MΦs, in human skin *in vivo*, were round and moderately dendritic (Fig. 4d), and they had a higher TPE-AF intensity *in vivo* compared to the ECM, as previously reported *in vitro*(Malissen et al., 2014; Njoroge et al., 2001).

**Figure 4.**
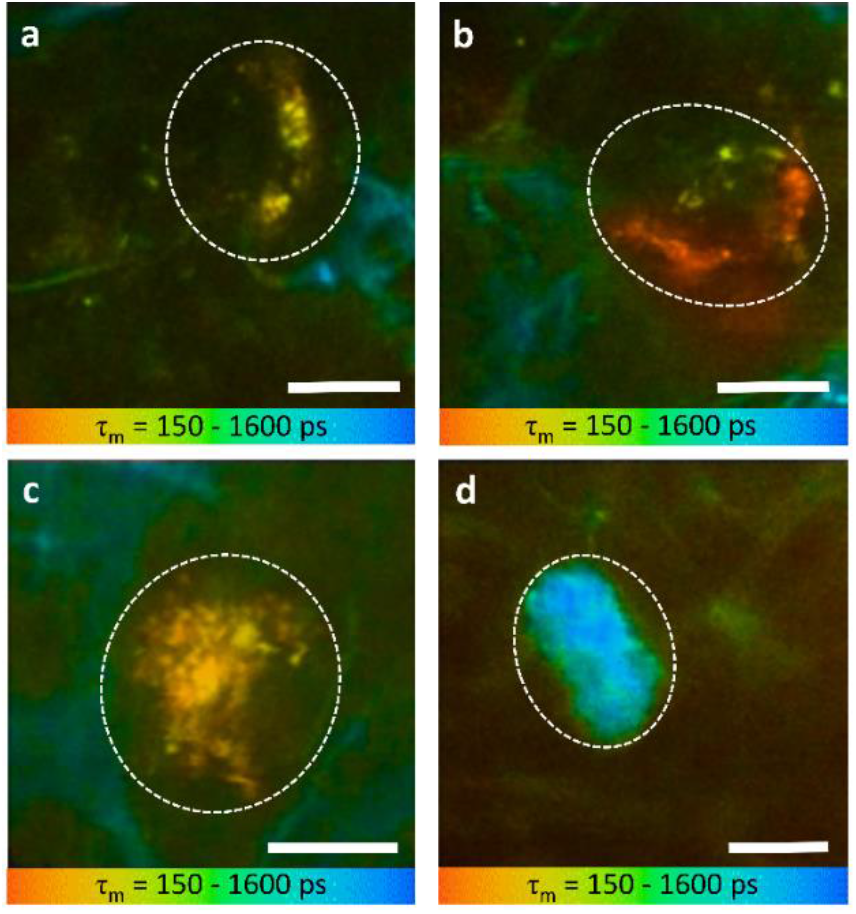
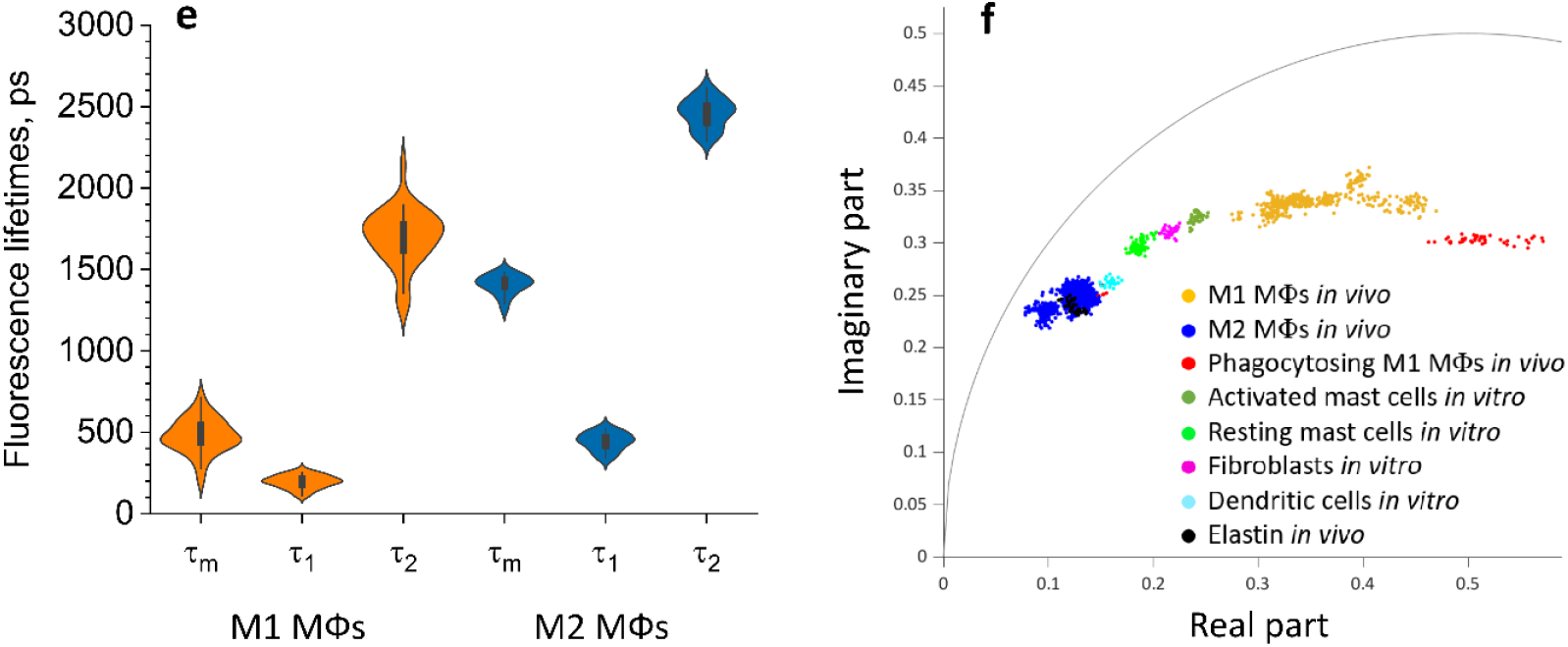
MΦs are visualised and categorised by TPE-FLIM signatures *in vivo*. TPE-FLIM *in vivo* images of potential perivascular flat spindle shaped M1 MΦ (a), of suspected slightly dendritic M1 MΦ in the depth 90 μm (b) large intervascular M1 MΦ with membrane extensions (c) and *in vivo* dermal cells resembling M2 MΦ were observed with a significantly longer mean fluorescence lifetime *τ*m compared to M1 MΦs and less pronounced TPE-AF intensity (d), showing mean fluorescence lifetime *τ_m_* in colour gradient from 150 to 1,600 ps. Scale bar: 10 μm. The histogram shows the distribution of TPE-FLIM parameters for *in vivo* MΦs (e). The boxplot represents 25–75% of the values. The phasor plot has a threshold at 0.9 of the maximum intensity and shows a summary of 12 M1, 2 phagocytosing M1 MΦs and 12 M2 MΦs *in vivo*, (f), where M1 MΦs are in orange and M2 MΦs in blue and phagocytosing M1 MΦs in red, the other dermal components are shown from *in vitro* measurements. The *in vivo* images (a–d) were recorded at 760 nm excitation wavelength, 50 mW laser power and 6.8 s acquisition time, in the depth of 80–100 μm on the volar forearm skin area of 25 healthy human subjects.

The TPE-FLIM parameters of *in vivo* M1 MΦs were in agreement with those of *in vitro* monocyte-derived and dermal M1 MΦs and *ex vivo* M1 MΦs (Fig. 4e, Table 1). M2 MΦs *in vivo* have longer *τ*_m_ fluorescence lifetimes compared to *in vitro* and *ex vivo* experiments. Yet, the *τ*_1_ and *τ*_2_ were in agreement with *in vitro* PBMC-derived monocytes, and the size and morphological parameters were in line with what is expected in M2 MΦs. The 2D segmentation in Fig. S5 shows the distinction of M1 and M2 MΦs presented in Figs. 4a-d, and the phasor plot in Fig. 4f shows that M1 and M2 MΦs could be distinguished from each other and from other dermal cells and ECM.

### TPE-FLIM can distinguish resting from phagocytosing human skin M1 MΦs in vivo

Phagocytosing skin M1 MΦs are characterized by an increase in cell size(May and Machesky, 2001), enhanced vacuolization(Cheng et al., 2019) and a shift of TPE-FLIM parameters towards shorter fluorescence lifetime values(Yakimov et al., 2019), different from those of resting M1 and M2 MΦs. A dermal cell matching these criteria is visualized *in vivo* using TPE-FLIM and presented in Fig. 5. This cell is located in the reticular dermis, has an enlarged size (≈25 μm) and an oval shape, similar to the resting M1 MΦ in Fig. 4c, pronounced vacuole structure and short TPE-FLIM lifetime specific for phagocytosing M1 MΦ. Of 37 dermal M1 MΦs analysed, 2 showed phagocytosis, and both were located in the reticular dermis below 100 μm of depth.

**Figure 5.**
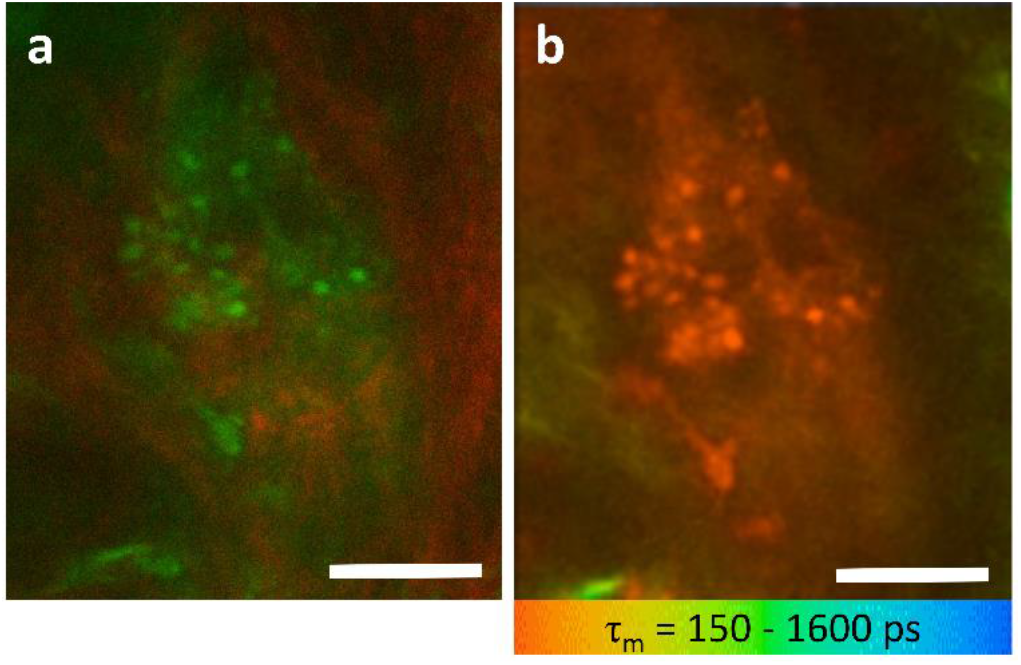
TPE-FLIM allows for visualization of phagocytosing M1 MΦs *in vivo*. Merged TPE-AF (green) and SHG (red) image of activated (phagocytosing) M1 MΦ measured *in vivo* in human papillary dermis at the depth 105 μm (a) and corresponding TPE-FLIM image (b), showing mean fluorescence lifetime *τ_m_* in colour gradient from 150 to 1,600 ps. Scale bar: 10 μm. The images were recorded at 760 nm excitation wavelength, 50 mW laser power and 6.8 s acquisition time on the plantar forearm.

### Classification algorithm to identify MΦs in the skin

To separate M1 and M2 MΦs from other dermal cells, we developed a classification algorithm, which used the decision tree (Fig. S6) and automatically classified MΦs based on their TPE-FLIM parameters and morphological features. The parameters of the decision tree were improved using hyperparameter optimization. The splitting method in the nodes of the decision tree classifier is chosen to be entropy impurity. To ensure the optimal quality of a split in the node of the decision tree, the following requirements had to be fulfilled: The minimal samples for a split is 2, the maximum depth of the tree is 9, and the samples had equal weight for the model classifying M1 and M2 MΦs. The ground truth was established by classification of *in vitro* and *ex vivo* MΦs with known phenotype resulting in 0.95±0.05 sensitivity and 0.97±0.06 specificity. When MΦs were classified as one group against other dermal cells, the sensitivity was 0.81±0.03 and the specificity was 0.81±0.03. Our algorithm also distinguished M1 MΦs from M2 MΦs and other cells, with a sensitivity of 0.88±0.04 and a specificity 0.89±0.03. For distinction of M2 MΦs from M1 MΦs and other cells, the sensitivity was 0.82±0.03 and the specificity was 0.90±0.03 (receiver operating characteristic is shown in Fig. S7.

## Discussion

This is the first *in vivo* study to show that human skin MΦs can be distinguished from other dermal cells and quantified through visualization with label-free, completely non-invasive TPE-FLIM. This risk-free approach also allows for the identification of MΦ phenotypes, i.e. M1 and M2 MΦs, and for the characterization of their functional stage, i.e. resting vs phagocytosing M1 MΦs. Finally, TPE-FLIM can be used to implement sensitive and specific machine learning algorithms for MΦ detection in the skin.

Our initial work with CD-14 positive monocytes isolated from PBMC and then differentiated and polarised towards M1 (IFN-γ) and M2 (IL-4) MΦs was needed to establish their TPE-FLIM parameters. In fact, it showed that MΦs are fluorescence-active, and, more importantly, that their TPE-FLIM parameters are among the best differentiators of M1 (*τ_m_*=479±106) and M2 (*τ_m_*=1,185±170) MΦs. M1 MΦs are associated with a slightly higher TPE-AF intensity (Table 1), which is a prominent indicator for the metabolic stress of the cell on account of a shift in lifetimes by changing amounts of free and bound NAD(P)H (Al-Shabany et al., 2016) and generation of ROS in mitochondria, phagosomal vacuoles and the cell membrane (Datta et al., 2015). Additionally to NAD(P)H, autofluorescence of lipids and other cell compartments was recorded. TPE-AF intensity is a parameter with limitation due to the nonlinear imaging technique. There is no linear correlation between excitation and emission intensity, also it is reduced due to scattering and absorption in the skin. The longer fluorescence lifetime *τ_2_* in M2 MΦs is best explained by oxidative phosphorylation and the emergence of fluorophores caused by fatty acid oxidation(Viola et al., 2019). NAD(P)H fluorescence is ubiquitously present in cells and exhibits the continuum of lifetimes in the 360–3,400 ps range. Free NAD(P)H has a short lifetime of 360 ps. For bound NAD(P)H, longer lifetimes up to 2–4 ns have been reported(Alfonso-García et al., 2016). A higher ratio of bound to free NAD(P)H is associated with M2 MΦs resulting in longer TPE-FLIM parameters, while a lower ratio of bound to free NAD(P)H is associated with M1 MΦs resulting in faster TPE-AF decay(Blacker et al., 2014). Thus, a strong indicator for the MΦ polarization is the TPE-FLIM parameters of monocytes in between cohorts of MΦs.

Translation of our *in vitro* findings to MΦs isolated from human skin confirmed that the latter share the TPE-FLIM signatures of the former, with shorter *τ_m_* in dermal M1 MΦs and longer *τ_m_* in dermal M2 MΦs. The classification into M1 and M2 MΦs *in vitro* based on their distinct TPE-FLIM parameters was supported by their differences in size, morphology and internal vacuole structure. That the *τ_1_* lifetime of dermal M2 MΦs is longer than that of monocyte-derived M2 MΦs is most likely due to the use of different polarization agents. MΦ colony-stimulating factor (M-CSF) and IFNγ for M1 MΦs and MΦ colony-stimulating factor (M-CSF) and IL-4 for M2 MΦs were used in PBMC-derived MΦs and microenvironment effects like inflammatory signals, UV exposure (Kang et al., 1994) and immune responses (Theret et al., 2019) influencing MΦ functions, result in divergent fluorescence lifetimes (Zhang et al., 2014) (Table 1, Fig. 2d). The most important outcome of our work with dermal MΦs was the establishment of their phenotype-specific TPE-FLIM signatures, a prerequisite for our subsequent *in vivo* studies and for comparing skin MΦs and other dermal cells.

In fact, the use of the TPE-FLIM signatures of M1 and M2 MΦs clearly allowed to distinguish them from mast cells, dendritic cells, fibroblasts and neutrophils (Table 1, Fig. 4f) (Kröger et al., 2020). We controlled for this by independent markers. For example, MΦs exhibited a higher fluorescence intensity compared to other dermal cells. In addition, they were larger than dermal mast cells, and their morphology was clearly different from that of neutrophils and fibroblasts. Vacuoles, a defining feature of MΦs, were linked to MΦ TPE-FLIM parameters, whereas granules, which identify mast cells, were not. In short, dermal MΦs, in their TPE-FLIM profiles, do not superimpose with other dermal cells or structures. The only exception, a partial overlap between M2 MΦs and elastin, is not relevant for the visualization of the former, as they are readily distinguished from the latter based on their morphology.

To confirm that *ex vivo* TPE-FLIM overlap with TPE-FLIM signatures of M1 and M2 MΦs *in vitro*, we sequentially analyzed cells in skin biopsy cryosections with TPE-FLIM and conventional immunohistochemistry. Indeed, both approaches identified and distinguished matching MΦ populations, i.e. M1 and M2 MΦs, with strong fluorescence intensity and spindle shape appearance of M1 MΦs and lower fluorescence intensity and longer fluorescence decay in M2 MΦs (Fig. 3). Interestingly, M2 MΦs are often found in an area of higher density of unknown dermal cells, presumably fibroblasts, compared to M1 MΦs. It is suspected that M2 MΦs in conjunction with collagen synthesizing fibroblasts are acting towards and aiding in dermal repair and regeneration. However, this approach also revealed some challenges that come with MΦ visualization by TPE-FLIM. For example, it was more difficult to visualise M2 MΦs than M1 MΦs in biopsies due to the high fluorescence intensity of elastin in dried tissue and other ECM components and a decreased signal-to-noise ratio. In Fig. 3d more CD163 positive M2 MΦs are visible compared to the corresponding TPE-FLIM image in Fig. 3c, which is due to previously mentioned challenges and the limited imaging plane of the two-photon tomograph (1.2–2.0 μm) compared to significantly thicker biopsy section (10 μm), which was stained in an entire depth and visualized by bright field microscopy. Importantly, immunohistochemistry confirmed our MΦ phenotype-specific TPE-FLIM signatures, and, in addition, confirmed that they distinguish MΦs from other dermal cells. Skin mast cells, for example, stained for tryptase, showed a distinct TPE-FLIM signature, confirming our recently reported findings on dermal mast cells *in vivo* (Kröger et al., 2020), and distinguished them from M1 and M2 MΦs (Fig. S8).

When we turned to the visualization of skin MΦs *in vivo*, we had to first develop a search strategy. Important considerations included the preferred localization of MΦs in the papillary and reticular dermis, the orientation of the focal plane parallel to the skin surface, and the need for maximal cellular cross-section visualization, which requires a high-resolution adjustment in depth to reconstruct an entire cell structure. The application of this search algorithm successfully visualised M1 and M2 MΦs in human skin *in vivo*. The *in vivo* TPE-FLIM parameters of M1 MΦs were in agreement with those observed *in vitro* and *ex vivo*. M2 MΦs *in vivo* were characterised by longer mean fluorescence lifetime *τ*m compared to *in vitro* and *ex vivo* (Table 1), which can be explained by the influence of environment (Koo and Garg, 2019; Njoroge et al., 2001). MΦs measured *in vivo* differ from cells measured *in vitro* by their simplified microenvironment (Mosser and Edwards, 2008) with missing growth factors and cytokines (Melton et al., 2015) and an elevated level of nutrients, which leads to different polarization of MΦs and different contributions of fluorescence lifetimes. Membrane extensions were harder to detect *in vivo* due to the obscuring effect of the surrounding ECM. We also observed that TPE-FLIM parameters in the same MΦs can vary depending on their cellular substructures, e.g. the nucleus, vacuoles, cytoplasm, or membrane (Figs. 4a–d). The phasor plot shows the relative position of the categories of MΦs and other cells. Furthermore, it shows the contributions of long and short fluorescence lifetimes and their discrepancies (Table 1) are due to the computational method and the harmonics at the repetition frequency of 80 MHz. Further investigations are needed to clarify how the location, morphology, and function of M1 and M2 MΦs influence their TPE-FLIM parameters and TPE-AF intensity *in vivo*. Such studies should also investigate the reasons for the differences in TPE-FLIM parameters between M1 and M2 MΦs, which may include differences in their metabolic pathways. M1 MΦs, for example, rely on NADH oxidase and production of ROS, which is linked to short fluorescence lifetimes of under 250 ps and mitochondrial fission. M2 MΦs, on the other hand, rely on oxidative phosphorylation and fatty acid oxidation, together with mitochondrial fusion (Ramond et al., 2019; Swindle et al., 2002; Xu et al., 2016).

The ability to visualise M1 and M2 MΦs by TPE-FLIM *in vivo* also makes it possible to explore how and why MΦs’ morphology, location and functions are linked. When activated, the cytoskeletal structure and cellular appearance of MΦs change, and this may also affect their TPE-FLIM parameters. M1 MΦs are elongated, with a dense actin network along the cortex. M2 MΦs are more spherical with more randomly distributed actin (Porcheray et al., 2005; Vogel et al., 2014; zhang et al., 2014) (Figs. 4b and 4d). Actin reorganization in M1 and M2 MΦ polarization and activation lead to bigger filament bundles of the actin cytoskeleton, which reduces cell plasticity (Colin-York et al., 2019; Pergola et al., 2017). As reported by Daphne and coworkers (Vogel et al., 2014), MΦ migration in the skin depends on their polarization. M1 MΦs, due to changes in actin cytoskeleton, migrate less far than M2 MΦs. Our TPE-FLIM findings confirm this, as we detected M1 MΦs via their high fluorescence and short autofluorescence lifetimes primarily in close proximity to blood capillaries. The irregular appearance of M1 MΦ detected by TPE-FLIM is likely a consequence of polarization-specific changes of the cellular cytoskeleton (McWhorter et al., 2013). The only morphological feature observed in both, M1 and M2 populations of MΦs, is that they are moderately dendritic, possibly because such MΦs are in the process of polarization, prior to cytoskeletal changes (Sica and Mantovani, 2012), or because polarization in MΦs is reversible polarizing (Sica and Mantovani, 2012; Yuan et al., 2017). Future studies should characterise the influence of cytoskeletal changes on TPE-FLIM parameters in detail and use TPE-FLIM to assess the impact of age, gender, and disease on the ratio, localization and function of M1 and M2 MΦs in the skin (Fukui et al., 2018).

During phagocytosis, the generation of ROS by NAD(P)H oxidase leads to the highest degree of metabolic stress observed in M1 MΦs besides apoptosis (Dupré-Crochet et al., 2013; Shirshin et al., 2019), and ROS localization in vacuoles in phagocytosing M1 MΦs as a bactericidal mechanism (Dupré-Crochet et al., 2013; Myers et al., 2003). This is why phagocytosing M1 MΦs change their TPE-FLIM lifetimes towards shorter values and their vacuoles become visible as localised bright spots, which makes their *in vivo* detection possible (Fig. 5b, Table 1) (Cannon and Swanson, 1992). TPE-FLIM allowed for the detection of phagocytosing M1 MΦs but cannot detect what they phagocytised. No internal structure was visible in TPE-FLIM images.

The construction of the feature vector and the resulting hyperparameter optimised decision tree model (Fig. S6) yielded proficient results for the automatised classification of M1 and M2 MΦs, demonstrating that M1 and M2 MΦs can be separated from each other and other cells in the skin with high accuracy, i.e. sensitivity and specificity, without need of additional staining using a supervised machine learning approach. The accuracy of this approach can be improved further, by the introduction of a depth-adjusted cell size and refined cell shape parameters and by increasing the number of *in vivo* MΦs integrated into the algorithm and training data set.

## Materials and Methods

### Two-photon excited fluorescence lifetime imaging (TPE-FLIM)

For imaging of human MΦs, a two-photon tomograph (Dermainspect, JenLab GmbH, Jena, Germany), equipped with a tuneable femtosecond Ti:sapphire laser (Mai Tai XF, Spectra Physics, USA, 710–920 nm, 100 fs pulses at a repetition rate of 80 MHz), was used at 3–5 mW for measurements of cells *in vitro* and skin biopsy sections *ex vivo*, as well as human dermis *in vivo* at 40–50 mW. The excitation wavelength was set to 760 nm, and a 410–680 nm band pass filter was used to detect two-photon excited autofluorescence (TPE-AF), whereas a 375–385 nm band pass filter was used to detect the second harmonic generation (SHG) signals. The axial and lateral resolution was approximately 1.2–2.0 and 0.5 μm, respectively(Breunig et al., 2013). The penetration depth covers the entire papillary dermis (Darvin et al., 2014; König, 2008; Kröger et al., 2020).

Fluorescence decay of a specimen was recorded and analysed in the SPCImage 8.0 software (Becker&Hickl, Berlin, Germany). TPE-FLIM data were fitted with a bi-exponential decay function. The TPE-AF intensity threshold was chosen depending on the signal-to-noise ratio, minimizing noise in the region of interest. The shift of the signal in relation to the instrument response function (IRF) was compensated. The typical IRF value was <100 ps. The TPE-AF decay curves were averaged over the central pixel of the region of interest and the 48 closest square neighbouring pixels (binning = 3), resulting in a number of detected photons for each fluorescence decay curve larger than 5,000. The TPE-AF decay parameters, decay lifetimes (*τ_1_* and *τ_2_*) and amplitudes (*a_1_* and *a_2_*), were used for the evaluation of the fluorescence lifetime distributions and 2D segmentation(Shirshin et al., 2017). The analysed parameters were the mean lifetime, defined as *τ_m_*=(*a_1_τ_1_*+*a_2_τ_2_*)/(*a_1_*+*a_2_*) and the ratios *τ_2_/τ_1_*, *a_1_/a_2_* and (*a_1_*-*a_2_*)/(*a_1_+α_2_*), which were used for 2D segmentation analysis. The TPE-FLIM data were also analysed and represented as phasor plots, that are based on the transformation of the fluorescence decay data in the frequency domain, whereas the decay is described as amplitude and phase values of the first Fourier component(Digman et al., 2008). The phasor plots’ *x*-axis is described by the cosine of the phase value multiplied by the amplitude, the *y*-axis represents the sine of the phase value multiplied with the amplitude(Lakner et al., 2017; Shirshin et al., 2019). The position of the mean lifetime is on the secant from *τ_1_* and *τ_2_*, the distance to the circle is given by the proportion of *a_1_* and *a_2_*. The TPE-FLIM data were normalised to the maximum intensity and the threshold of 70% was set when analysing the phasor plots.

### Ethical considerations and study conduct

Volunteers for intravital imaging provided written informed consent before participation. Skin samples taken from periocular skin surgery for MΦ preparation and all human skin investigated in this study were used after written informed consent was obtained. Positive votes for the experiments have been obtained from the ethics committee of the Charité – Universitätsmedizin Berlin (EA1/078/18, EA4/193/18, EA1/141/12), which were conducted according to the Declaration of Helsinki (59th WMA General Assembly, Seoul, October 2008).

### Study subjects

25 healthy volunteers (12 males and 13 females, 24–65 years old, skin type I–III according to Fitzpatrick classification(Fitzpatrick, 1988)) with asymptomatic volar forearm skin without preexisting health conditions were randomly selected for non-invasive *in vivo* measurements in the papillary dermis using TPE-FLIM. Visually impairing hair was removed with a scissor prior to measurements. The oil immersion objective of the microscope was connected to the skin via a 150 μm thick, 18 mm diameter cover glass (VWR, Darmstadt, Germany) with a ≈10 μL distilled water droplet between cover glass and skin. The volunteers were screened between October 2018 and November 2020.

### Investigation of human dermal MΦs in vitro

Human dermal MΦs were prepared from periocular tissue(Botting et al., 2017). Human periocular skin was digested in 2.4 U/ml dispase type II (Roche, Basel, Switzerland) at 4 °C for 12 hours. The dermis was minced with scissors after removal of the epidermis and further digested in PBS containing Ca^2+^ and Mg^2+^ (Gibco, Carlsbad, CA, USA) supplemented with 1% Pen/Strep, 5% FCS, 5 mM MgSO_4_, 10 μg/ml DNaseI (Roche, Basel, Switzerland), 2.5 μg/ml amphothericin (Biochrom, Berlin, Germany,) 1.5 mg/ml collagenase type II (Worthington Biochemical Corp., Lakewood, NJ, USA) and 0.75 mg/ml H-3506 hyaluronidase (Sigma-Aldrich, St. Louis, MO, USA) at 37 °C in a water bath with agitation for 60 min. The cell suspension was filtered using 300 μm and 40 μm stainless steel sieves (Retsch, Haan, Germany). Centrifugation at 300 × g for 15 min at 4 °C was applied next. The digestion cycle was repeated once. MΦs were isolated by Pan Monocyte Isolation Kit (Miltenyi, Bergisch Gladbach Germany) after washing in PBS w/o Ca^2+^ and Mg^2+^ (Gibco, Carlsbad, CA, USA), and kept in basal Iscove’s medium supplemented with 1% Pen/Strep, 10% FCS, 1% non-essential amino acids, 226 μM *α*-monothioglycerol (all Gibco, Carlsbad, CA, USA). For long-term cultures, after 24 h recombinant human IL-4 (20 ng/ml) and hSCF (100 ng/ml) (both Peprotech, Rocky Hill, NJ, USA) were added. Purity of MΦ cultures was routinely checked to be >85%(Nielsen et al., 2020). For imaging, cells were used after 3 days in medium, washed twice with PBS before seeding on 18 mm diameter microscope cover glass (VWR, Darmstadt, Germany) for imaging in PBS containing Ca^2+^ (Gibco, Carlsbad, CA, USA) at room temperature.

### Investigation of peripheral blood monocytes in vitro

Peripheral blood monocytes were isolated from human blood using 15 Ml Ficoll-Paque (VWR, Darmstadt, Germany) centrifugation gradient. Centrifugation was performed at 1,000 × g for 1 minute, with added 9 M1 heparin and filled to 50 Ml with PBS. Centrifugation was then repeated at 1,000 g for 10 minutes, discarding the upper plasma layer and collecting the PBMC layer. The cells were washed twice with PBS and centrifuged at 350 × g for 10 minutes. The supernatant was discarded and cultured in 5 M1 basal Iscove’s medium supplemented with 1% Pen/Strep, 10% FCS (Biochtrom, Berlin, Germany) and subsequently incubated at 37 °C and 5% CO_2_ for 2 h before seeded and imaged on an 18 mm diameter microscope cover glass (VWR, Darmstadt, Germany) in PBS containing Ca^2+^ (Gibco, Carlsbad, CA, USA) at room temperature.

### Investigation of MΦs differentiated from peripheral blood monocytes in vitro

MΦs were differentiated from peripheral blood monocytes and polarised into M1 (IFNγ)-like state with MΦ colony-stimulating factor (M-CSF) and IFNγ and M2 (IL-4)-like state with MΦ colony-stimulating factor (M-CSF) and IL-4. For further stimulation, cells were incubated with LPS at 37 °C for 24 hours prior to imaging.

### Investigation of dendritic cells in vitro

CD14 positive peripheral blood mononuclear cells were used to differentiate dendritic cells by washing in PBS and centrifuging at 350 × g for 10 minutes twice. 5 Ml RPMI medium, supplemented with 1% Pen/Strep and 1% FCS (Biochtrom, Berlin, Germany), was added. Tryptan Blue (Sigma-Aldrich, St. Louis, MO, USA) was used for counting the cells in a hemocytometer, seeded at 2.0 × 10^6^ cells/ml and incubated for 2 hours at 37 °C under 5% CO_2_. Non-attached cells and the supernatant were discarded. Adding 500 μl basal Iscove’s medium to the cells supplemented with 1% Pen/Strep, 1% glutamine, 5% HSA, (all Gibco, Carlsbad, CA, USA) 100 ng/ml IL-4, 100 ng/ml GM-CSF (both Peprotech, Rocky Hill, NJ, USA) with medium change every second day for 6 days at 37 °C. For TPE-FLIM imaging the cells were seeded on 18 mm diameter microscope cover glass (VWR, Darmstadt, Germany) in PBS containing Ca^2+^ at room temperature.

### Preparation and cryo-sectioning of human skin for combined TPE-FLIM and histomorphometric analysis

Thirteen human skin biopsy cryo-sections were prepared and measured using the TPE-FLIM method to acquire TPE-FLIM parameters of suspected M1 and M2 MΦs. The skin biopsies were obtained from abdominal reduction surgery of four female patients (31, 33, 40 and 44 y. o., skin type II according to Fitzpatrick classification(Fitzpatrick, 1988)). Punch biopsies of 6 mm diameter were obtained, frozen and stored at −80 °C before cryo-sectioning. Vertical histological cryo-sections of 10 μm thickness were prepared on a cryostat (Microm Cryo-Star HM 560, MICROM International GmbH, Walldorf, Germany) after embedding in a cryo-medium (Tissue Freezing Medium, Leica Biosystems Richmond Inc., Richmond, IL, USA) and placed on 18 mm diameter microscope cover glasses (VWR, Darmstadt, Germany). The anatomical condition of the biopsies was continuously examined using a transmission microscope (Olympus IX 50, Olympus K.K., Shinjuku, Tokyo, Japan).

Using TPE-FLIM, cryo-sections were searched for cells with MΦ-specific TPE-FLIM parameters and the corresponding TPE-FLIM images of suspected MΦs were recorded. To prove the measured cells are MΦs, the skin biopsies were labeled by irradiating a squared area of 28 μm×28 μm located near the suspected MΦs with a Ti:sapphire laser (Mai Tai XF, Spectra Physics, USA, 100 fs pulses at a repetition rate of 80 MHz) at a maximal power of 50 mW at 760 nm for 3 seconds. All incubations were performed at room temperature unless otherwise stated. In brief, sections were fixed for 10 min in cold acetone (−20 °C) and rinsed in TBS (Agilent, Santa Clara, CA, USA). For staining of MΦs, the MΦ-specific anti-CD68 (clone ab955) (Abcam, Cambridge, UK), Recombinant Anti-CD163 antibody [EPR14643-36] (clone ab189915) (Abcam, Cambridge, UK) were used to account for M1 and M2 MΦ phenotypes, respectively. Slides were rinsed three times with TBS, and endogenous peroxidase was blocked with 3% H_2_O_2_ in TBS for 5 min followed by incubation with anti-mouse EnVision+ labelled polymer (Agilent Technologies, Santa Clara, USA) for 30 min. Slides were rinsed in TBS as before and incubated with AEC substrate-chromogen (Agilent Technologies, Santa Clara, USA) for 10 min. Nuclei were counterstained with Mayer’s hemalum solution (Merck, Darmstadt, Germany). Stained MΦs have a brown-red color, which enables to visually distinguish them from other cells and the ECM. After the staining procedure, target MΦs and squared labels of the skin sections were identified by light microscopy and overlaid with TPE-FLIM images matching an appropriate magnification and image orientations.

Specifically, CD68-stained M1 MΦs were counted in the papillary dermis region in each biopsy, and an average of 209±25 cells/mm2 for the papillary dermis and an average of 140±76 cells/mm2 for the reticular dermis for a 10 μm deep cryo-section was observed (Fig. 1c). The density of the CD163 stained M2 MΦs was an average of 242±126 cells/mm2 for the papillary dermis and an average of 107±60 cells/mm2 for the reticular dermis for a 10 μm deep cryo-section (Fig. 1d).

The MΦs search algorithm we then used was similar to that recently presented by our group for the identification of resting and activated mast cells in the papillary dermis(Kröger et al., 2020) and included the following steps: First, the papillary dermis (≈60–100 μm depth for volar forearm) was explored for fluorescent spots of 10–15 μm in size with irregular shape and a membrane extension having bright spots of about 1–3 μm. The TPE-FLIM parameters of the suspected bright areas were measured and matched those of M1 and M2 MΦs obtained *in vitro* and *ex vivo*.

To prove that the TPE-FLIM parameters of other dermal cells, which have detectable TPE-AF intensity, namely, mast cells and dendritic cells do not match or superimpose with TPE-FLIM parameters of MΦs, negative control measurements were performed. The procedure was similar as described for the verification of MΦs in skin biopsies using specific immunofluorescence, but six human skin cryo-sections were stained for the presence of mast cells and two for dendritic cells.

Staining of mast cells was done by blocking with serum-free protein followed by incubation for 1 h with anti-tryptase antibody (clone AA1) diluted 1:1,000 in antibody diluent (all Agilent, Santa Clara, CA, USA). For staining of dendritic cells, anti-CD11c antibody (clone B-Ly6) (BD Biosciences, Franklin Lakes, NJ, USA) was used after fixing the cryo-section for 10 min in cold acetone (−20 °C) and rinsing with TBS.

### Statistical analysis and classification algorithm

Matlab R2016a (MathWorks, Natick, MA, USA) was applied for descriptive statistics of all TPE-FLIM data. All results are indicated as mean±standard deviation. Differences between distributions were compared using the non-parametric Kolmogorov-Smirnov test with a significance level of *α*=0.05. The decision tree classifier was modelled using Scikit-learn 0.22 in a Python 3.7 environment (Python Software Foundation, Delaware, USA). A randomised training set, consisting of 50% of the complete data set, was used for training and validating the test set 10,000 times. The true positive- and true negative rates were calculated from the confusion matrix and describe the quality of the classification and indicate type I and type II errors. For the decision tree(Breiman et al., 1984), the TPE-FLIM parameters *τ_1_*, *τ_2_*, *τ_m_*, *τ_1_/τ_2_*, *a_1_*, *a_2_*, *a_1_/a_2_*, (*a_1_*-*a_2_*)/(*a_1_*+*a_2_*), TPE-AF intensity, cell shape, and decay curve were used for each cell measured *in vitro*, *ex vivo*, and *in* and hyperparametrically optimized(Yang and Shami, 2020). The feature vector was constructed as follows: 1 (shape (circular form)) + 1 (normalised TPE-AF intensity of cell normalised by the cell’s area and excitation power) + 256 (fluorescence decay curve) + 8 (TPE-FLIM parameters obtained after bi-exponential approximation of the decay curve), in total, 266 values. The MΦ size was not included in the feature vector for the classification model, as MΦs *in vivo* could have slightly different dimensions from those measured *in vitro* (in cell cultures) and *ex vivo* (in biopsies), caused by obscuring effects of surrounding dermal tissue. Here, 1 represents circular and 0 non-circular shape. The lifetimes calculated from the biexponential decay model were averaged over the whole cell, and the fluorescence intensity was normalised by optical power and averaged the pixel of interest and the 48 neighbouring square pixel.

In total, 110 MΦs *in vitro*, 20 MΦs *ex vivo*, 70 MΦs *in vivo* (for M1/M2 ratio see Table 1), 59 mast cells *in vitro*, 17 mast cells *ex vivo*, 82 mast cells *in vivo*, 14 dendritic cells *in vitro*, 6 fibroblasts *in vitro* and 21 neutrophils *in vitro* were used as input for the model (399 cells in total). Given data vectors from *x_i_* ∈ *R^n^*, *i*=1,…,*l* and a label vector *y_i_* ∈ *R^l^*, where a decision tree recursively separates the data into two classes with the mode *m* represented as *Q*. For each node a split *θ* = (*j, t_m_*) decided with the feature *j* and the threshold *t_m_*. The node split the data into subsets *Q_left_*(*θ*) and *Q_right_*(*θ*)

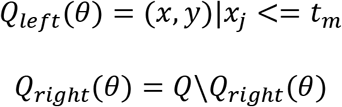

The impurity was calculated by the impurity function *H()* at the mode *m*

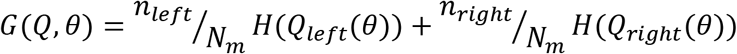

With the parameters for minimised impurities the subsets were recourse until *N_m_* =1.

The returns values of the classification were 0 for M1 MΦs, 1 for M2 MΦs and 2 for other dermal cells, 0 for MΦs and 1 for other dermal cells for node *m* in the region *R_m_* and *N_m_* observation, the proportion of class *k* observations in node *m* is *p_mk_* = *1/N_m_* ∑_*x_i_*∈*R*_*m*__ *I*(*y_i_* = *k*).

Receiver operating characteristic (ROC)-curves served as a tool to determine the diagnostic abilities of the method, where the true positive rate was plotted against the false positive rate of the respective outcomes for both the categorization of MΦs against other dermal cells and M1 MΦs and M2 MΦs against other dermal cells.

## Supplementary Materials

**Fig. S1.**
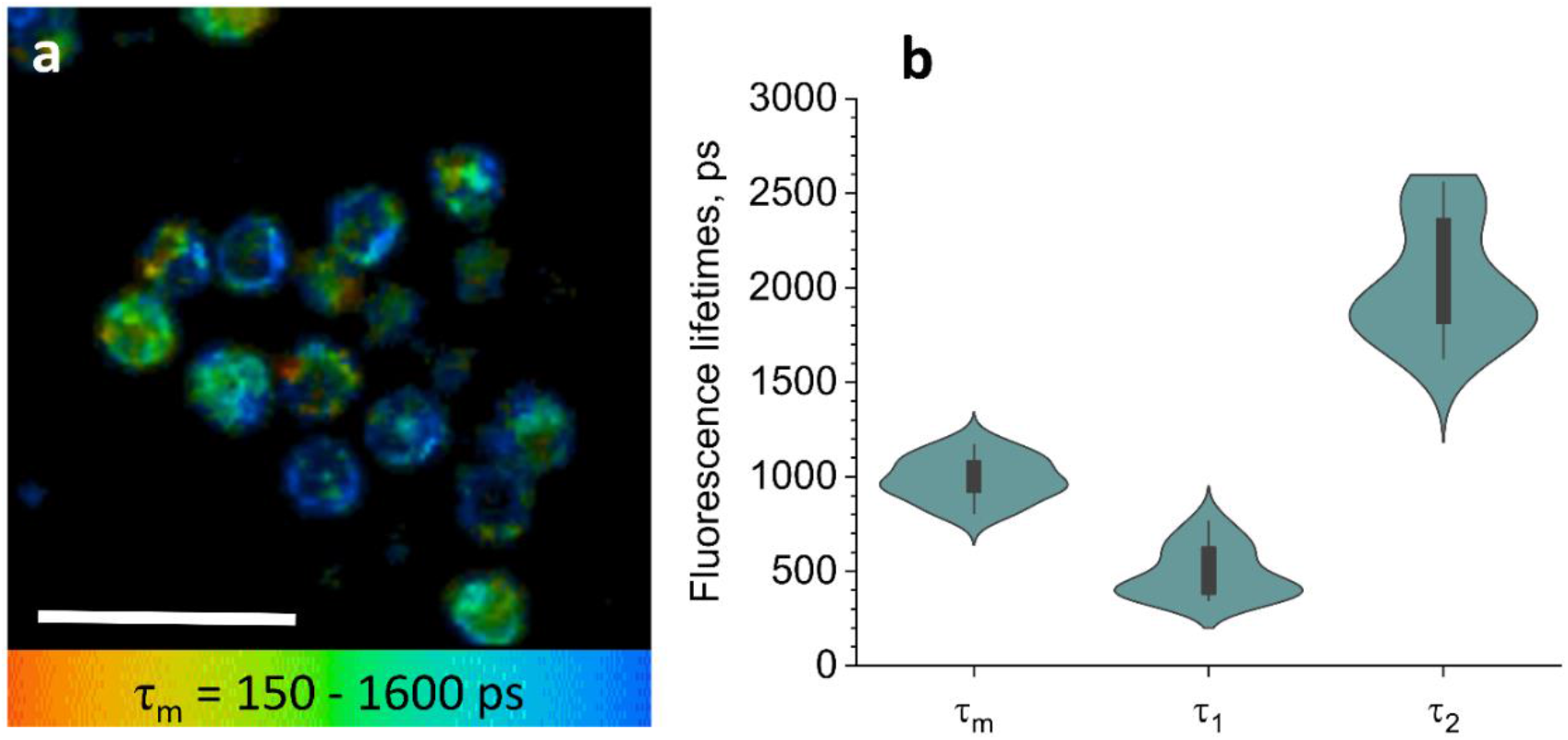
TPE-FLIM visualisation of PBMC and histogram of TPE-FLIM parameter. TPE-FLIM *τ_m_* images (mean fluorescence lifetime *τ_m_* in the 150–1,600 ps range) of PBMC (a) Scale bar: 10 μm. The distribution of TPE-FLIM parameters *τ_1_*, *τ_2_* and *τ_m_* for PBMC (*n*=15) (f).

**Fig. S2.**
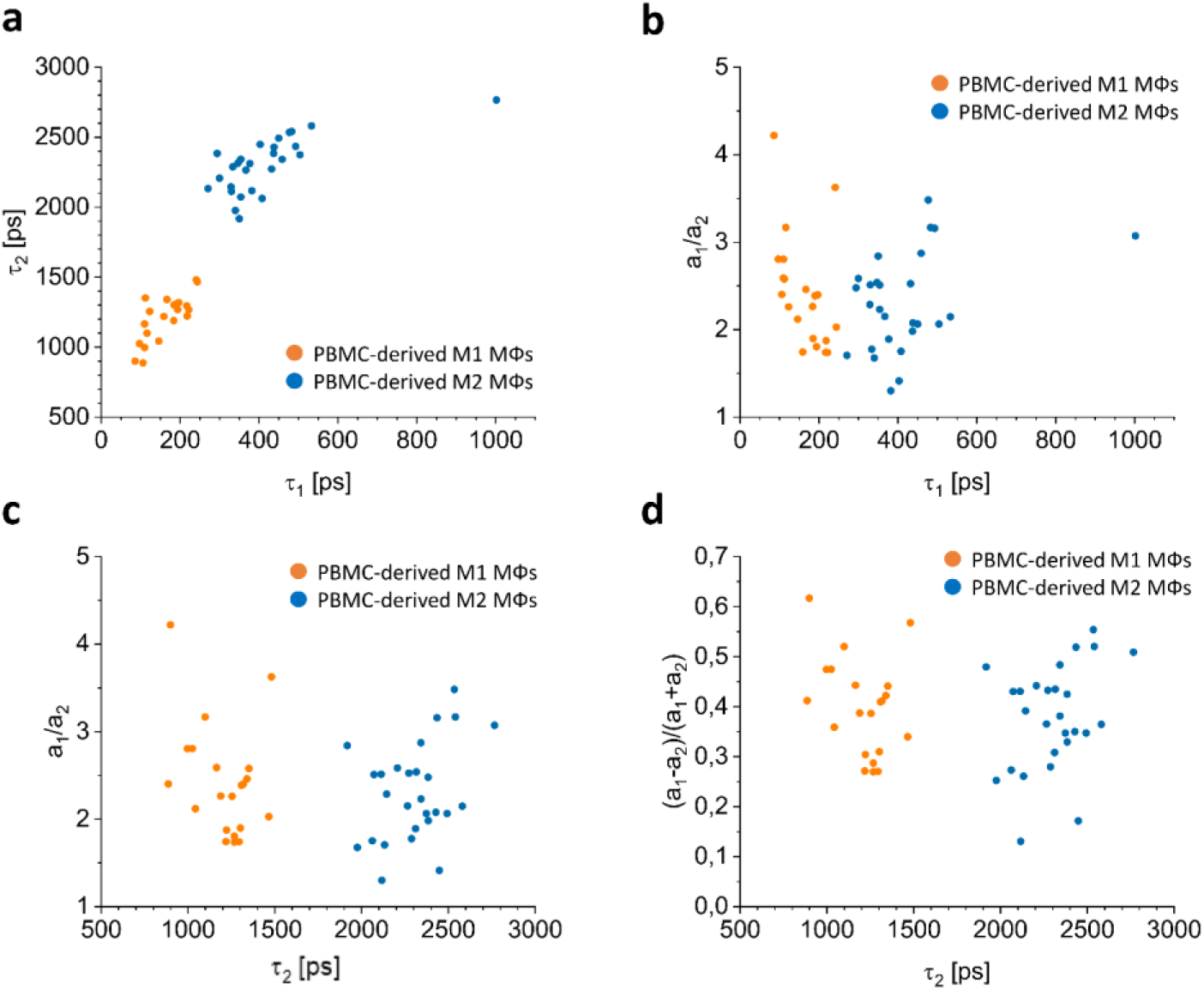
*In vitro* 2D segmentation for MΦs from PBMC. 2D segmentation of the *τ_1_*(*τ_2_*) (a), *τ_1_*(*a_1_*/*a_2_*) (b), *τ_2_*(*a_1_*/*a_2_*) (c) and *τ_2_*((*a_1_*-*a_2_*)/(*a_1_*+*a_2_*)) (d) TPE-FLIM parameters of the M1 MΦs (orange circles) and M2 MΦs (blue circles) measured *in vitro*.

**Fig. S3.**
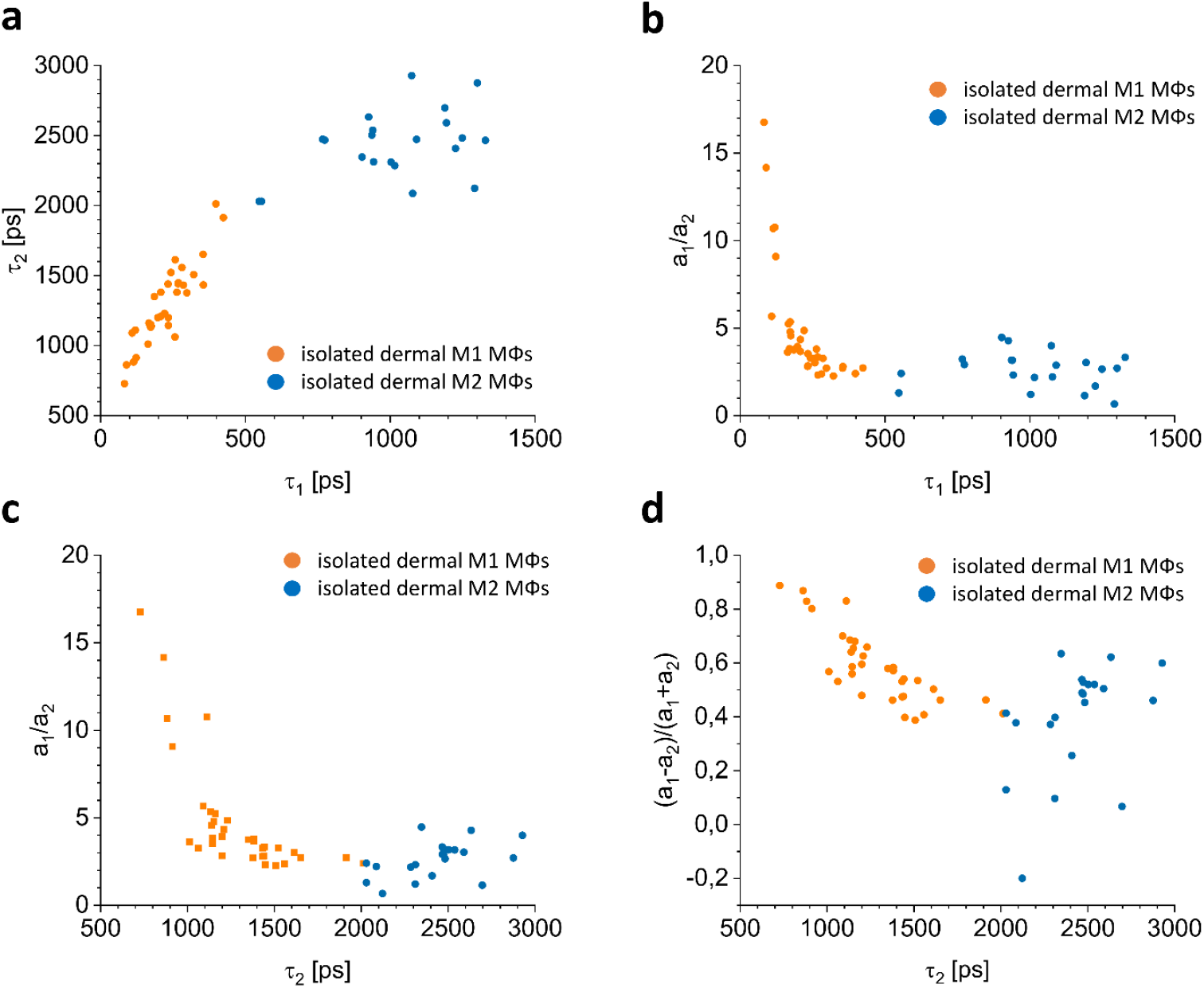
*In vitro* segmentation for MΦs from dermal tissue. 2D segmentation of the *τ_1_*(*τ_2_*) (a), *τ_1_*(*a_1_/a_2_*) (b), *τ_2_*(*a_1_*/*a_2_*) (c) and *τ_2_*((*a_1_*-*a_2_*)/(*a_1_*+*a_2_*)) (d) TPE-FLIM parameters of the M1 MΦs (orange circles) and M2 MΦs (blue circles) measured *in vitro*.

**Fig. S4.**
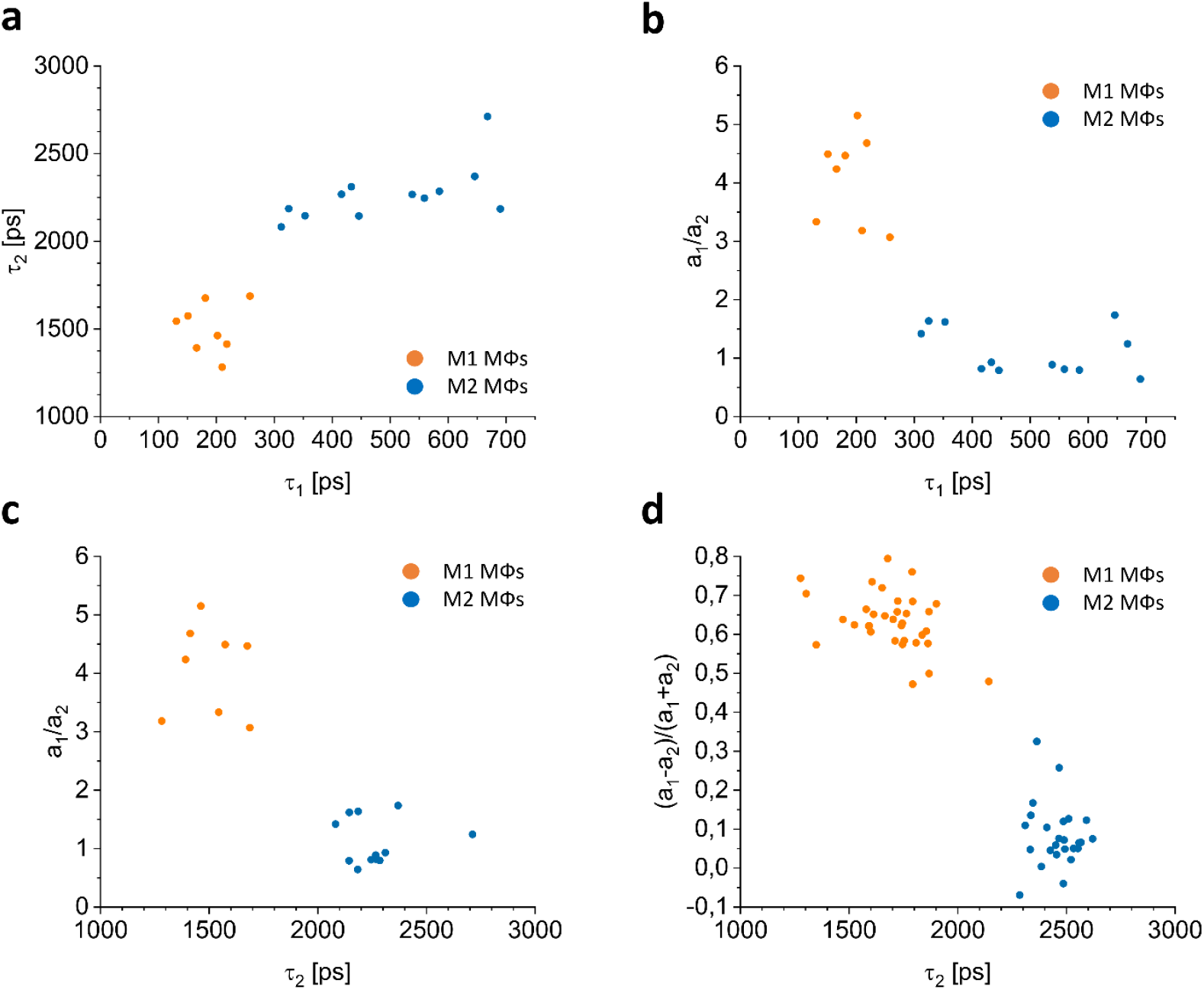
*Ex vivo* segmentation for MΦs from skin biopsies. 2D segmentation of the *τ_1_*(*τ_2_*) (a), *τ_1_*(*a_1_*/*α_2_*) (b), *τ_2_*(*a_1_*/*a_2_*) (c) and *τ_2_*((*a_1_*-*a_2_*)/(*a_1_*+*a_2_*)) (d) TPE-FLIM parameters of the M1 MΦs (orange circles) and M2 MΦs (blue circles) measured *ex vivo*.

**Fig. S5.**
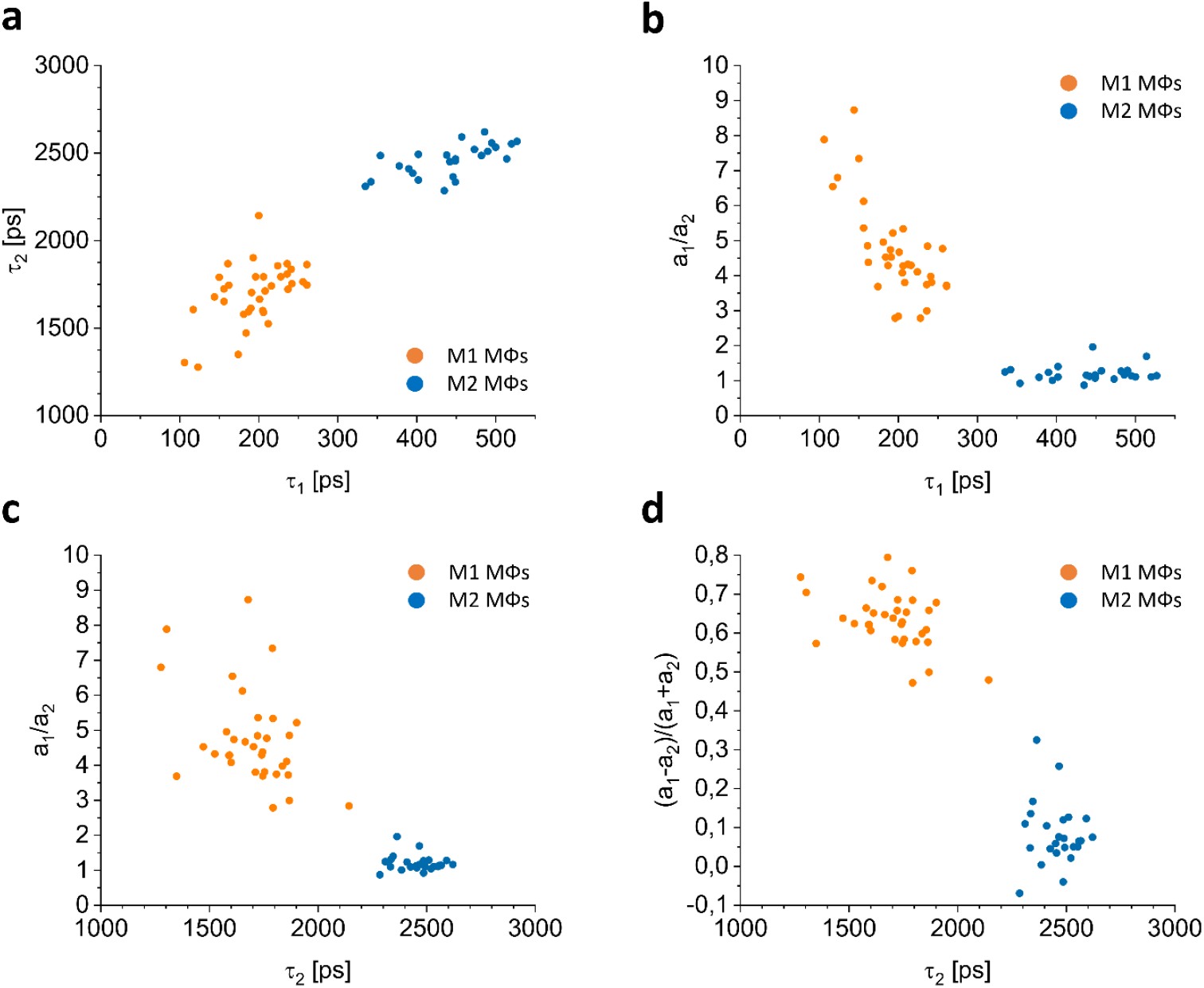
*In vivo* segmentation for MΦs in human skin. 2D segmentation of the *τ_1_*(*τ_2_*) (a), *τ_1_*(*a_1_/a_2_*) (b), *τ_2_*(*a_1_/a_2_*) (c) and *τ_2_*((*a_1_*-*a_2_*)/(*a_1_*+*a_2_*)) (d) TPE-FLIM parameters of the M1 MΦs (orange circles) and M2 MΦs (blue circles) measured *in vitro*.

**Fig. S6.**
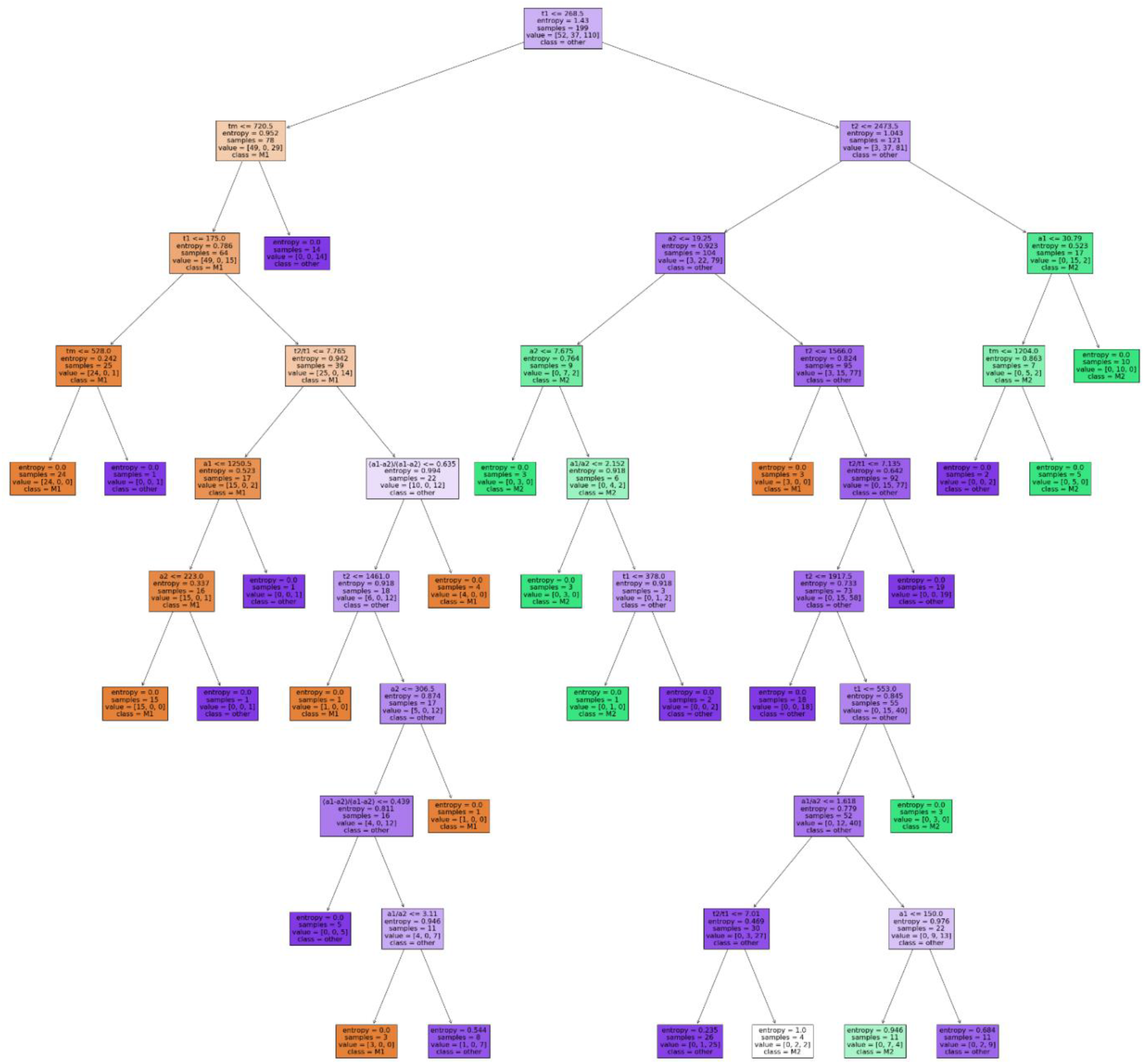
Decision tree model. Decision tree model for classification of M1 and M2 MΦs with the parameters entropy impurity, minimal samples for split is 2, maximum depth of the tree is 9, and the samples had equal weight. The text of the decision tree is readable when zoomed in.

**Fig. S7.**
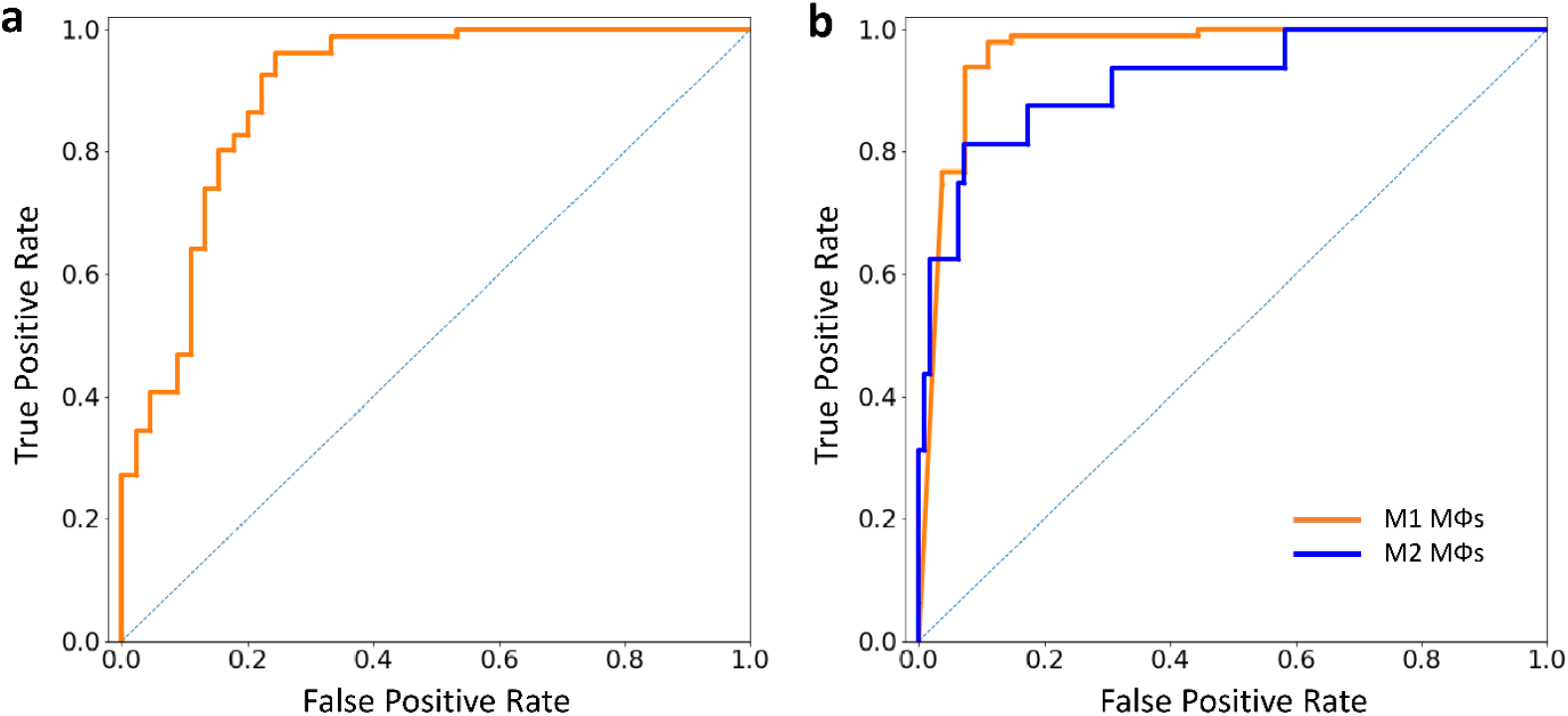
ROC curves of decision tree models. ROC curves of false positive to false negative rates for classification of MΦs versus other dermal cells (a) and for classification of M1 MΦs (orange line) versus M2 MΦs and other dermal cells and for classification of M2 MΦs (blue line) versus M1 MΦs and other dermal cells (b) generated from the decision tree classifier (Fig. S6).

**Fig. S8.**
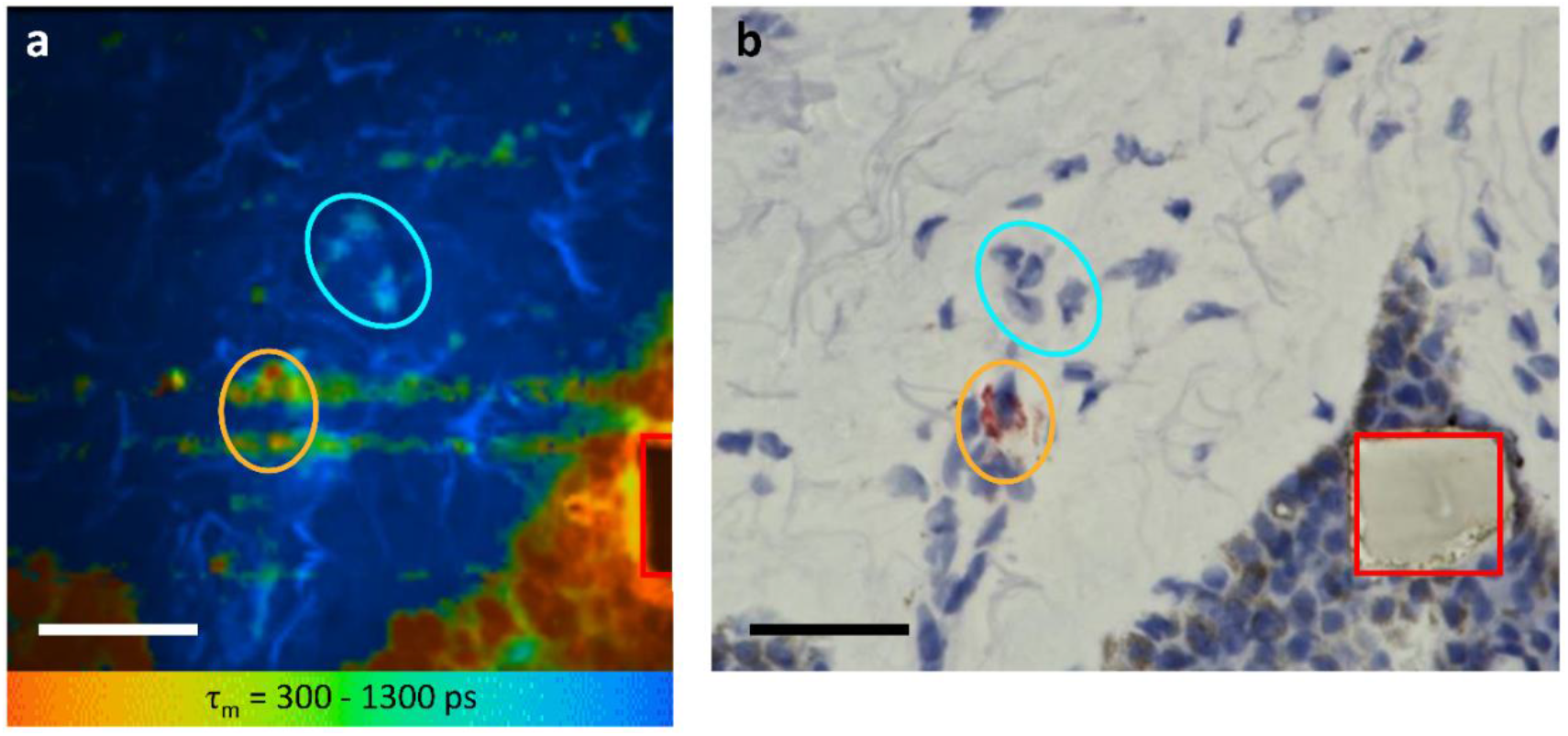
Mast cell-specific staining with tryptase *ex vivo* – negative control. Side by side comparison of TPE-FLIM image (average lifetime *τ_m_* in the 300–1,300 ps range) (a) and corresponding light microscopic image after staining of human skin biopsy cryo-sections for MCs (b). Laser power is 3 mW, excitation at 760 nm. The MCs are marked with orange ellipses, suspected MΦs are marked with blue ellipses, and the laser burned squares (28×28 μm^2^) used for labeling are marked in red. Images have been rotated and zoomed to match the same orientation and size. Scale bar: 30 μm.

## General

We thank Evelin Hagen, Niklas Mahnke and Loris Busch from Charité – Universitätsmedizin Berlin for their excellent technical support. We thank David Satzinger for the schematic illustration of MΦs.

## Funding

Foundation for Skin Physiology of the Donor Association for German Science and Humanities (MK, JS, MCM, JL, MED) Russian Science Foundation №19-75-10077 «Photonic and Quantum technologies. Digital medicine» (ES).

## Author contributions

M.E.D., J.L. and M.M. planned and designed the study. M.K., M.E.D., J.Sche., J.Schl., E.A.S., M.C.M. J.L. and M.M. conceived the experiments. M.K. and M.E.D. performed the experiments. M.K., J.Sche., J.Schl., E.A.S. and M.E.D. analyzed the data. M.E.D., M.K. and M.M. wrote the main manuscript text. All authors reviewed the manuscript.

## Competing interests

The authors declare no competing interests in relation to this work, or other interests that might be perceived to influence the results and/or discussion reported in this paper.

Marcus Maurer is or recently was a speaker and/or advisor for and/or has received research funding from Allakos, Amgen, Aralez, ArgenX, AstraZeneca, BioCryst, Celldex, Centogene, CSL Behring, FAES, Genentech, GIInnovation, Gilead, Innate Pharma, Kalvista, Kyowa Kirin, Leo Pharma, Lilly, Menarini, Moxie, Novartis, Pharming, Pharvaris, Roche, Sanofi/Regeneron, Shire/Takeda, Third HarmonicBio, UCB, and Uriach.

## Notes

https://datadryad.org/stash/share/H_R3MdaIi-ZsjNnmaQ_N3wkbc90dFn0ZSFUhE2CXjbc

### Competing Interest Statement

The authors have declared no competing interest.

